# Estimation of mutual information for real-valued data with error bars and controlled bias

**DOI:** 10.1101/589929

**Authors:** Caroline M. Holmes, Ilya Nemenman

## Abstract

Estimation of mutual information between (multidimensional) real-valued variables is used in analysis of complex systems, biological systems, and recently also quantum systems. This estimation is a hard problem, and universally good estimators provably do not exist. Kraskov et al. (PRE, 2004) introduced a successful mutual information estimation approach based on the statistics of distances between neighboring data points, which empirically works for a wide class of underlying probability distributions. Here we improve this estimator by (i) expanding its range of applicability, and by providing (ii) a self-consistent way of verifying the absence of bias, (iii) a method for estimation of its variance, and (iv) a criterion for choosing the values of the free parameter of the estimator. We demonstrate the performance of our estimator on synthetic data sets, as well as on neurophysiological and systems biology data sets.

## I. INTRODUCTION

Much of 20th century statistical physics was built by studying dependences among physical variables expressed through their variances and covariances. However, in recent decades, physicists have started to explore systems (particularly those far from equilibrium), where correlation functions, which are the most useful in the context of small fluctuations and perturbative calculations, do not tell the whole story about the underlying systems, which exhibit large, nonlinear fluctuations. A related problem is that correlation functions depend on the choice of a parameterization used to measure observables, so that, for example, for large fluctuations, the correlation between *x* and *y* can be very different from that between log *x* and log *y*, making it harder to interpret the data.

A common solution to these problems is to use the mutual information between two variables instead of their correlation to quantify dependence [1, 2]. Mutual information between variables *x* and *y* is distributed according to a joint distribution *P* (*x, y*) is defined as

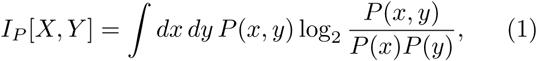

where the integral should be interpreted as a sum for discrete variables, and as a multi-dimensional integral for multi-dimensional real-valued variables. Mutual information quantifies *all*, and not just linear dependences between the two variables: it is zero if and only if the variables are completely statistically independent [2]. Further, mutual information does not change under invertible transformations (reparameterizations) of *x* and *y* [2]. These properties make mutual information the quantity of choice for analysis of dependences between real-valued, nonlinearly-related variables, especially in modern biophysics (see Refs. [3–5] for just a few examples).

An important complication that prevents an even wider adoption of information-based analyses is that mutual information and related quantities are notoriously difficult to estimate from empirical data. Mutual information involves averages of logarithms of *P*, the underlying probability distribution.Since, for small *P*, – log_2_ *P* → ∞, the ranges of *x, y* where *P* is small and hence cannot be sampled and estimated reliably from data contribute disproportionately to the value of information. In other words, unlike correlation functions, information depends nonlinearly on *P*, so that these sampling errors result in a strong sample size dependent and *P* -dependent bias in information estimates. In fact, even for discrete data, there can be no universally unbiased estimators of information until the number of samples, *N*, is much larger than the cardinality of the underlying distribution, *K* [6]. This means that, for continuous variables, universally unbiased information estimators do not exist at all. These simple observations have resulted in a lively field of developing entropy / information estimators for discrete variables, which work under a variety of different assumptions (see [6–13]). Such estimators often use one of the following ideas. First, for *N* ≫ 1, when most possible outcomes have been observed in the sampled data, one may hope that the bias of an estimator can be written as a power series in 1*/N*, and then the first few terms of the series can be calculated analytically, or estimated directly from data by varying the size of the data set. Second, coincidences start happening in data at much smaller *N* than it takes to sample every possible outcome [14]. One can then use the statistics of such frequently occurring outcomes to extrapolate and learn properties of the large low-probability tail of the distribution *P*, estimating contributions of the tail to the information. Third, one can estimate the bias of an estimator by applying it to a shuffled data set, where the mutual information is zero by construction. Some of these ideas can be applied to continuous variables as well, by soft or hard discretization of the data.

However, for many experiments dealing with continuous variables, such as when studying motor control, some of these bias correction approaches are not easily applicable [15, 16]. First, the observed variables may be very large dimensional, which makes good sampling nearly impossible. Second, when focusing on mutual information between just two variables that are projections of very large dimensional variables, shuffling may not work as a way to check bias. Indeed, for any finite *N*, shuffling is not guaranteed to remove statistical dependences among *all* data dimensions simultaneously, and randomizing along one set of projections may leave residual mutual information due to statistical dependences along the others. Thus developing information estimators that use continuity of real-valued data to help with undersampling, estimate information without resampling, and work for large-dimensional data is crucial. One of the most successful such estimators was proposed by Kraskov, Stögbauer, and Grassberger [17], which we will refer to as KSG. It uses distances to the *k*-th nearest neighbors of points in the data set to detect structures in the underlying probability distribution. If some points cluster, then the *x* coordinate of a point can be used to predict its *y* coordinate, resulting in a nonzero mutual information. This can be detected by the statistics of the *k*-th nearest neighbor distances. Further, by varying *k*, one can vary the spatial scale on which structures are detected.

While successful, KSG cannot be a universally good for all underlying probability distributions. In fact, even the original Ref. [17] pointed out that there are probability distributions for which the estimator does not converge to the right answer even at very large *N*. However, we are not aware of any published methods for self-consistently detecting if the estimator is unbiased on specific datasets. Our goal here is to make KSG more broadly useful by endowing it with the abilities (i) to estimate its own error bars, (ii) to detect existence of a sample-size dependent bias, and (iii) to automatically choose the hyperparameter *k* most appropriate for the current data. Further, (iv) we directly expand the range of probability distributions, for which the estimator remains unbiased, by using the reparameterization invariance property of the mutual information.

Some of the methods presented in this paper were first tried in Ref. [16], but here we test them more thoroughly, introduce additional changes, and formalize the approach. We start this paper with a brief review of the KSG estimator. We then progressively introduce our modifications of the method. Finally we give examples of performance of the modified method on simulated and real-life data sets.

### A. The KSG estimator

Mutual information can be written down as the difference of marginal and joint Shannon entropies [2]:

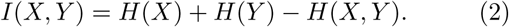

KSG uses the Kozachenko-Leonenko (KL) *k*th nearest neighbor entropy estimator [18] for each one of the differential entropy terms:

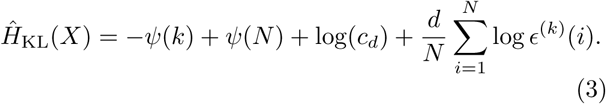

Here *ψ* is the digamma function, *d* is the dimensionality of *x, N* is the total number of samples, *c*_*d*_ is the volume of a unit ball with *d* dimensions, and *ϵ*^(*k*)^(*i*) is twice the distance between the *i*’th data point and its *k*’th neighbor. The intuition is that, if the distances *ϵ*^(*k*)^(*i*) are small, then the underlying probability distribution is concentrated, and the corresponding differential entropy is also small. Notice that the metric for calculating distances has to be defined *a priori* to apply this estimator, and the metrics can be very different in the *x* and the *y* spaces.

One could plug in Eq. (3) for each one of the three differential entropies in Eq. (2), but then the biases in the estimates of the marginal and the joint entropies likely will not cancel – if the ball with the radius *ϵ*^(*k*)^(*i*) includes the *k*th nearest neighbor of the *i*th data point in the *d*(*x*) + *d*(*y*) dimensional space, then the ball of the same radius will include a lot more data points in just *d*(*x*) or *d*(*y*) dimensions. Reference [17] argued that keeping the ball size rather than *k* constant for the marginal and the joint entropy would result in the decrease of the total mutual information bias. To implement this, KSG uses the max(Δ*x*, Δ*y*) metric to define the distance between two points that are (Δ*x*, Δ*y*) away from each other. It then defines the smallest rectangle in the (*x, y*) space centered at a point *i* that contains *k* of its neighboring points. One then denotes by 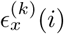 and 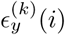 the *x* and *y* extents of this rectangle, and by 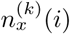 and 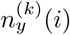 the number of points such that ‖*x*_*j*_ *– x*_*i*_‖ *≤ ϵ*_*x*_(*i*)*/*2 or ‖*y*_*j*_ *- y*_*i*_‖ *≤ ϵ*_*y*_(*i*)*/*2, respectively. Then the mutual information is estimated as [17, 19]

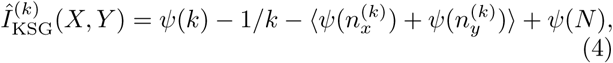

where averaging is over the samples. Note that, if 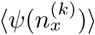 and 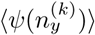 increase, the mutual information estimate drops. This can be understood intuitively as follows. First, recall that *ψ*(*n*) *→* log *n* for large values of the argument, and thus grows with *n*. Since *ψ*(*n*) is convex up, 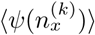 is large when 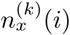 are narrowly distributed (and the same for *y*, respectively). But if values of 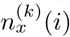 (or 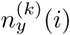) are nearly the same for all *i*s, then the underlying probability distribution has no structural features in the *x* (or *y*) direction, and the mutual information must be low, which is exactly what Eq. (4) suggests.

Empirically, KSG is one of the best performing mutual information estimators for continuous data. It has been used widely, with over 1700 citations to the original article according to Google Scholar as of the writing of this article. And yet some basic questions remain unanswered. Foremost is that *k* is a free parameter, which needs to be chosen before applying the estimator to data. Varying *k* allows one to explore features in the probability distribution across different spatial scales, resulting in the usual bias-variance tradeoff. For example, *k* = 1 will pick up even very fine features, but at the same time 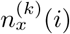 and 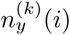 will be small, resulting in large fluctuations. On the other hand, large *k* may miss fine-scale features and hence underestimate the information, but statistical fluctuations will be smaller. One can expect that the optimal value of *k* depends on the structure of the spatial features in the data, which may be nontrivial and may exist on multiple spatial scales. In addition, the optimal *k* should also depend on *N*, since fine features can only be observed at high sampling density. Thus choosing the best *k* is not a simple task. The original KSG analysis focused largely on *N* → ∞ and on probability distributions with large, uniform spatial features, for which *k ∼ N* was often useful (though *k* = 2 … 4, which is small but not 1, was also recommended). In contrast, real life problems often have *N* ∼ 10^2^ … 10^4^ and many heterogeneous spatial features, so that only *k ∼* 1 may have a chance of working. In this article, in addition to other modifications, we propose a way of estimating an optimal value of *k* for KSG. Crucially, in order to do so, we first solve two other problems: estimating the standard error of the estimator and its bias directly from data.

## II. RESULTS

### A. Estimating the variance of KSG

We first focus on estimating the standard deviation of KSG. For this, we start with bivariate normally distributed data as a test case since, for such data, the choice of *k* has only a small effect on 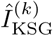 [17]. Additionally, for a bivariate Gaussian, the true value of mutual information is related to the correlation coefficient *ρ* as 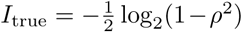, which allows for an easy determination of the actual error of the estimator. Specifically, for the rest of this section, we will frequently use *ρ* = 0.6 as an example, where *I*_true_ ≈ 0.32 bits.

For a single data set taken at random from this bivariate Gaussian, KSG will produce an estimate, e. g., 0.2802 bits for *N* = 1000. However, since we do not know the standard deviation of the estimator (its “error bars”), we do not know how many of these digits are significant, and whether the estimate is biased. Calculating the error bars is not simple since standard methods, such as bootstrapping, only work for quantities that are *linear* in the underlying probability distribution [20], while information is not. This is easy to understand intuitively: resampling data with replacements – a key step in bootstrap – creates duplicate data points. These will be interpreted by KSG as fine-scale, high-information features, leading to overestimation of the mutual information in the bootstrapped samples.

To illustrate the inadequacy of bootstrap for this problem, we generate 20 independent sets of data of size *N* = 200 from a bivariate Gaussian with *ρ* = 0.6. We then estimate 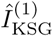 for each set, and finally calculate the mean and the standard deviation of these 20 KSG estimates. The result is 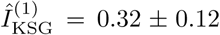, which matches well with the analytical value of ≈ 0.32 bits. On the other hand, if we take the single data set of *N* = 200 and then bootstrap it and calculate the mean and the standard deviation of the KSG estimates of the boot-strapped data, we get 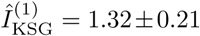. The mean is wrong by a factor of about 4, and even the standard deviation is twice as large as it should be (and the scale of both errors certainly depends on *N* and the underlying distribution). We emphasize this again: *bootstrapping, at least in its simple form, should not be used in estimation of mutual information or its error bars!*

Instead of using bootstrapping for estimating the error of KSG, we propose to use the fact that variance of essentially any function that, like Eq. (4), is an average of *N* random i. i. d. contributions scales as 1*/N* for sufficiently large *N*. Indeed, as seen in Fig. 1, this scaling holds, for example, for bivariate Gaussians with different correlation coefficients for, at least, *N* > 50.

**FIG. 1.**
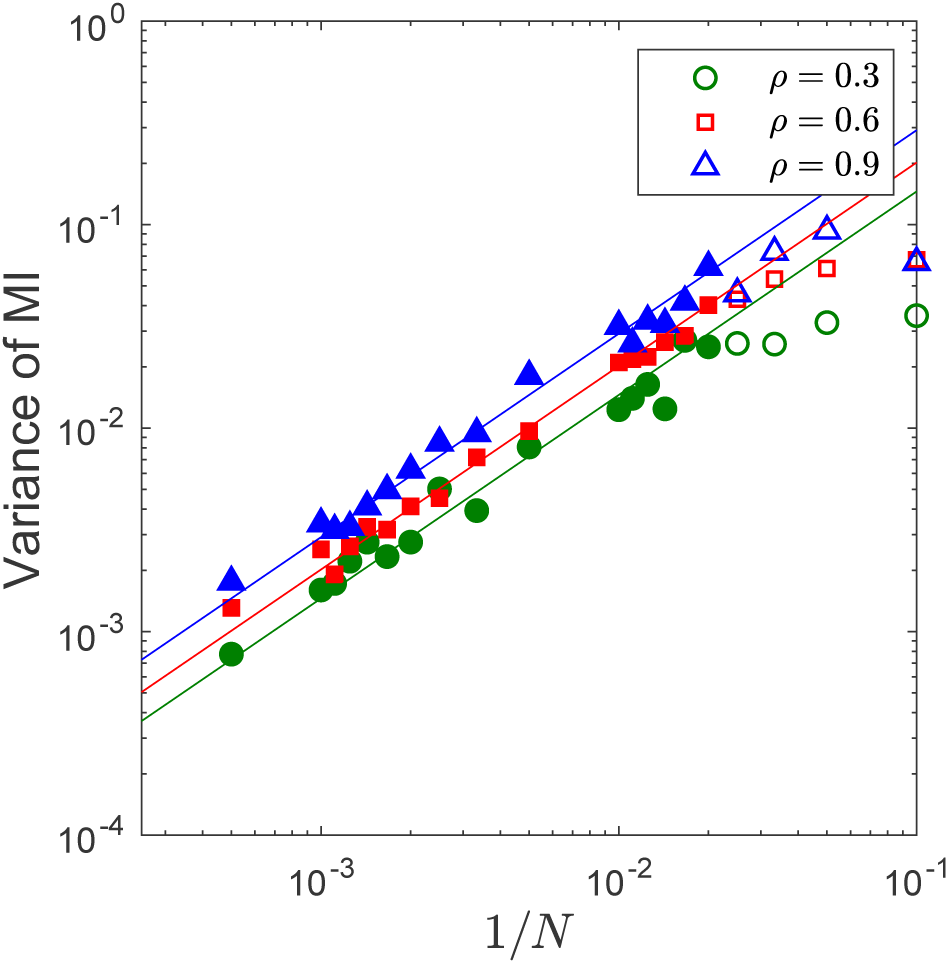
Dependence of the variance of KSG on the sample set size. For bivariate Gaussians with three different correlation coefficients *ρ* = 0.3, 0.6, 0.9, we generate 100 independent sample data sets of different sizes *N*. For each *N*, we calculate 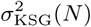 as the empirical variance of all 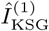 with this *N*. The variance is plotted vs. 1*/N*. The shown linear fit illustrates that the variance, indeed, scales as 1*/N* for *N* ≫ 1. Empty symbols were not used to fit the linear relation.

Thus we write for the variance of KSG

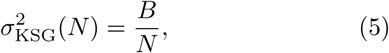

where the value of *B* will depend on the particular distribution. To estimate *B* for specific data, we subsample (*not* re-sample!) the data. Specifically, for a small integer *n*, we partition the data set of size *N* at random into *n* non-overlapping subsets of as close to equal sizes as possible. We calculate 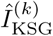 for each such subset. Then the sample variance of these *n* values of 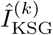 is our estimate of 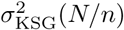. Once we know 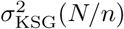 for many values of *n*, we fit the model, Eq. (5), to these values and estimate *B* empirically. Finally, knowing *B*, we calculate 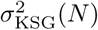 from Eq. (5) directly. Combining these steps, we get expressions for the estimate of the variance of the estimator, as well as the standard error of the variance itself, which can be found in the Appendix, Eqs. (9) and (10), respectively.

We finish the Section with a few observations. First, one might be tempted to generate many different non-overlapping partitions of the data at the same *n*, hoping to average over the partitions and hence decrease the variability observed in Fig. 2. This should be avoided since such different permutations of data would not produce independent samples of the variance. For the same reason, one should avoid any overlaps among partitions, so that the number of samples in each partition is *N/n* with an integer *n*. Finally, the 1*/N* scaling of the variance only works for large *N*. Thus it may not hold for *n* ≫ 1, limiting the maximum value of *n* in realistic applications. For all plots shown here, we use *n* = 1 … 10, which we generally find to be sufficient.

**FIG. 2.**
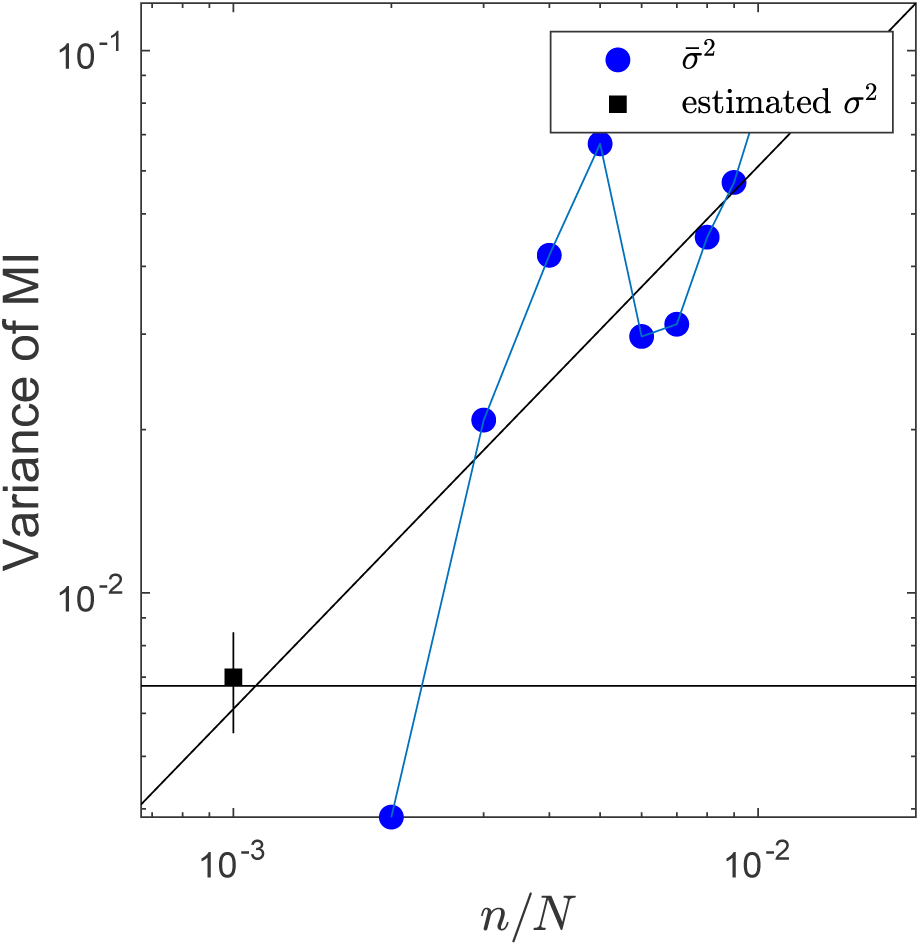
Calculating the variance of KSG. For a bivariate Gaussian with *ρ* = 0.6, we sample *N* = 1000 data points from the distribution. We calculate the variance of KSG with *k* = 1 for *N/n* data points by partitioning the data into *n* nonoverlapping subsets and estimating the mutual information for each subset, as described in the main text (blue dots). An unweighted linear fit with the slope of 1 is shown as a guide to eye, illustrating extrapolation of the variance of the estimator to the full data set size. An estimate of the variance of the estimator, with its own expected error, is performed using the analysis in the Appendix and is denoted by a black square with an error bar. For comparison, the horizontal line denotes the variance of the estimator calculated from applying it to 100,000 independent samples of size *N* = 1000 from the Gaussian, illustrating a near perfect agreement.

### B. Detecting the estimation bias and choosing *k*

Most common mutual information estimators, including KSG, are asymptotically unbiased for sufficiently regular probability distributions at *N* → ∞. At the same time, all are typically biased at finite *N*, as discussed in the Introduction. As a result, the bias is *sample size dependent*. Thus while it may be hard to calculate the bias analytically for specific data and estimators, one may be able to estimate it empirically by varying the size of the data set [8, 15, 16, 21]: if the estimated mutual information drifts with changing *N*, there are reasons to be concerned about the bias. Here we will use this strategy to ascertain the existence of a sample size dependent bias for KSG.

We note that, unlike Ref. [8], we are not interested in estimating the bias at finite *N* and then subtracting it out (equivalently, extrapolating 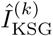 to *N* → ∞). This is possible only when the form of the bias as a function of *N* is known, leaving only a small number of coefficients to be characterized from data themselves, such as for the classical ∼ 1*/N* Miller-Madow correction to the maximum likelihood information estimator [22]. For KSG, the asymptotic scaling of the bias is unknown, making this approach currently infeasible. Further, any estimator would exhibit statistical fluctuations when applied to real data. Unless the standard deviation of the estimator is known, one cannot say whether the observed sample size dependent drift is due the bias or to the fluctuation: only if the systematic drift over a reasonable range of *N* is much larger than the standard deviation, would one consider this an evidence of the bias. Thus detecting the bias of KSG (or any other estimator) by varying *N* is impossible without a careful consideration of how 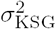 behaves.

The question of detecting the bias is intimately related to choosing *k*, the number of nearest neighbors considered by the estimator: we expect the bias to be *k*-dependent. Specifically, for large *k*, fine-scale features in the underlying probability distribution will be missed by KSG, and the mutual information will typically be under-estimated. At the same time, because 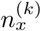 and 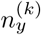 grow with *k*, we expect the standard deviation of the estimator to be smaller at larger *k*. In contrast, for smaller *k*, statistical fluctuations will be much larger, while two different effects will affect the bias. First, the downwards information bias is expected to be smaller at small *k* since finer scale features will be explored. Second, larger fluctuations in 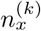 and 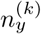 will lead to a larger *N* -dependent upwards bias in 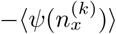 and 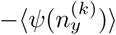 in Eq. (4). Overall, the bias at small *k* may be of an arbitrary sign. In any case, one can explore the drift as a function of *N* for different values of *k* and choose to work with the value (if one exists), for which (a) there is no sample-size dependent drift compared to the estimator standard deviation, and (b) the standard deviation is the smallest. We also note that the actual estimated value of the mutual information can be strongly *k*-dependent; we will discuss this further below, but we note here briefly that it is important for the estimated value of the information to be stable across a range of *k*’s.

We illustrate this analysis in Fig. 3 for the bi-variate normal distribution. Here we work with smaller data sets than in the previous figures to better explore the effects of *k*. Of the three values of *k* shown in the Figure, *k* = 4 shows the best combination of no sample size dependent drift and low variance. Correspondingly, as this drift analysis predicts, 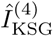 remains unbiased compared to the true mutual information value over the entire range of data explored. We also verified that the estimator is relatively stable to the choice of *k*, so that other values near *k* = 4 give similar 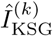, and the estimator remains unbiased (not shown).

**FIG. 3.**
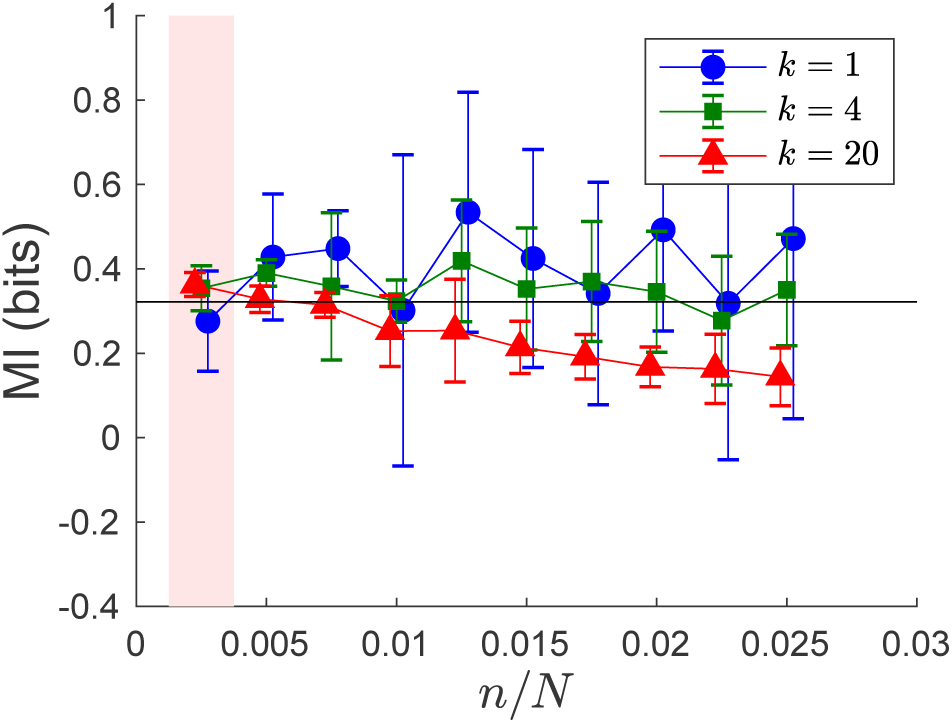
Bias of KSG as a function of *N* and *k*. Starting with *N* = 400 samples from a bivariate normal distribution with *ρ* = 0.6, we partition the data into *n* nonverlapping sub-samples (without replacements), each with *N/n* data points. We estimate 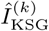 for each subsample using Eq. (4). Means and standard deviations of the estimates for each set of *n* partitions are shown for three different values of *k*. The leftmost point (on pink packground) for each line has error bars representing our estimate, following the methods we discussed in the previous section. The true mutual information of 0.322 bits is shown as a black horizontal line. For the data set sizes explored here, *N/n* = 40 … 400, *k* = 20 clearly leads to a statistically significant negative bias, while *k* = 1 gives an unnecessarily high variance, sometimes dipping into mathematically impossible negative values. *k* = 4 shows a low-bias, low-variance behavior for these *N/n*. Note that symbols for different values of *k* are slightly shifted relative to each other for visibility, but are actually evaluated at the same *n/N* for all *k*.

We note that Ref. [17] explored, in particular, *k* ∝ *N*, and *N* → ∞. In contrast, our approach often gives *k ∼* 1 for *N* ∼ 10^2^ … 10^4^. We expect that *k* ∝ *N*^*η*^ for some distribution-dependent *η <* 1 to be asymptotically optimal since it would lead to both (i) exploring progressively finer features and (ii) smaller relative fluctuations in 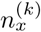 and 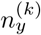 as *N* → ∞. However, here we are interested in applications to real experimental data sets. These are usually far from the asymptotic regime, so that the available range of *N* is too small to meaningfully think about different scalings of *k*.

### C. Decreasing the KSG bias

Empirically, KSG exhibits large biases for distributions that have very heavy tails, have structural features on multiple length scales, or are severely skewed. All of this can be traced to the non-symmetric distribution of data points in the *ϵ*-balls. As an example, Fig. 4 (A) shows application of KSG for different values of *k* to a bivariate log-normal distribution. Even for a very large *N* = 10000, KSG is severely negatively biased for all *k*s. In specific realizations, we often see the bias *increasing* as *N* grows, so that the KSG estimate turns negative, while mutual information must always be positive. We note that small negative values of information would not be a concern generally: in order to estimate information near zero bits with error bars, one needs to have it be negative sometimes — negative estimates that fall within error bars of zero are acceptable. Here, however, the estimates can be consistently and significantly negative, indicating a serious problem.

**FIG. 4.**
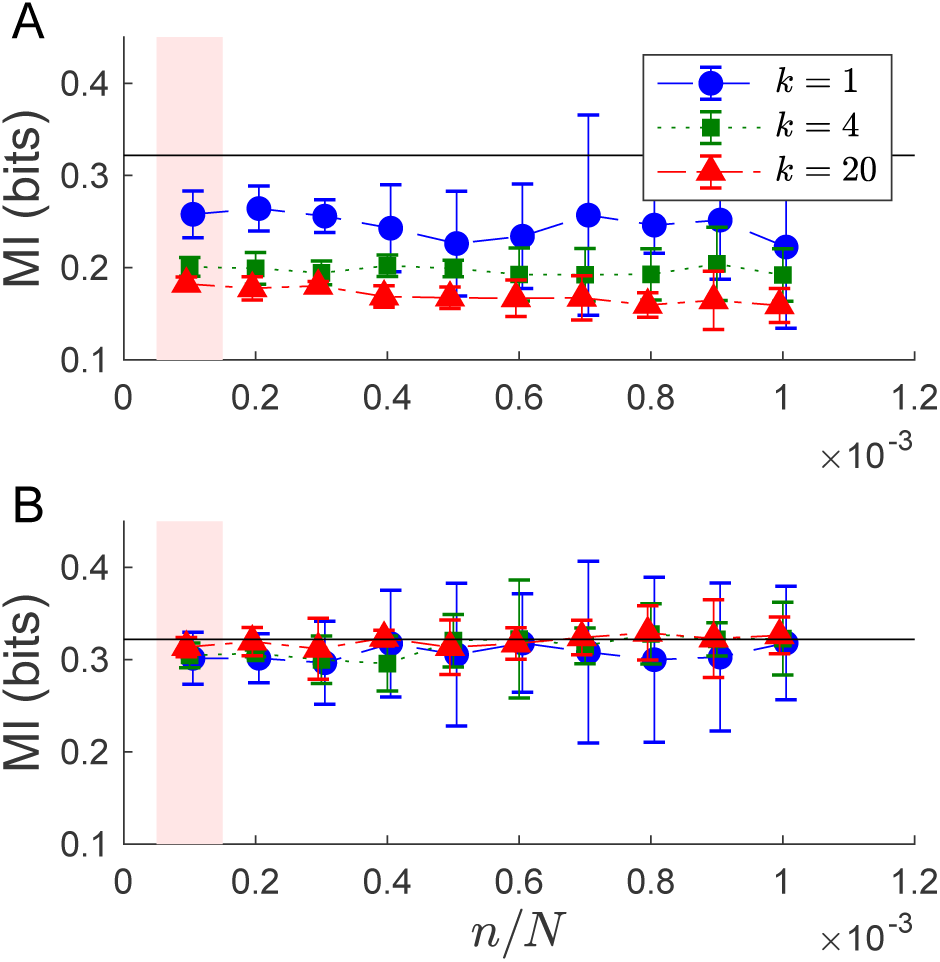
Marginally normalizing the data decreases the KSG bias. (A) For a bivariate log-normal distribution, *P* (*x, y*) ∼ exp (–((ln 3*x*)^2^ + (ln 5*y*)^2^ – 2*ρ* ln 3*x* ln 5*y*)*/*(2(1 – *ρ*^2^))), with *x* and *y* being standard normal, *ρ* = 0.6, and the true mutual information of 0.322 bits, we repeat the analysis from Fig. 3 and plot the dependence of 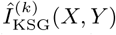 on *k* and *n/N* for *N* = 10^4^. As always, the error bars on the leftmost points (full data set, pink background) are estimated as discussed above. The true value of information is shown as a black horizontal line. KSG does not give a consistent estimate of the information, and any estimate would be a function of *k*. No value of *k* gives the correct mutual information. (B) After reparameterizing each marginal into a standard normal, we investigate the dependence of 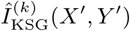 on *k* and *n/N*. Here KSG does not show a sample sign dependent drift and is, therefore, largely unbiased for all tested values of *k*. Here we also have an estimate that is independent of the choice of *k*.

However, as we mentioned above, mutual information is invariant under invertible marginal reparameterizations. Thus one can hope to increase the range of distributions for which KSG is unbiased, by reparameterizing the data to distributions that KSG is better equipped to handle. Specifically, since KSG works extremely well for normal variables [17], we suggest to transform each marginal variable *x* and *y* into a standard normal variable. For example, if we define *r*_*i*_ = 1 *… N* as the rank of the corresponding *x*_*i*_, then its reparameterized version is

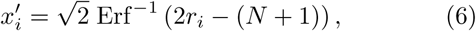

where Erf^*-*1^ is the inverse of the error function. Indeed, as illustrated in Fig. 4 (B), this transformation removes the bias for many cases. Note that we did not use the fact that the distribution is bivariate log-normal during the reparameterization: Eq. (6) will transform *any* data into marginally normal variables.

In some sense, the log-normal example is trivial, since marginal reparameterizations transform it not just into *marginally* normal, but into *jointly* normal distribution, which would not be expected generically.However, since KSG depends largely on *marginal* neighborhoods, cf. Eq. (4), one would expect that joint normality after reparameterization is not necessary, and marginal normality alone is sufficient for the bias to be decreased. Below we illustrate this on two real experimental datasets.

However, before that, we need first to show that our procedure for estimating the variance of the estimator can be used for reparameterized data, where biases may exist, and where the original distribution is non-gaussian. For this, we repeat the analysis of Fig. 1 for reparameterized data: Figure 5 shows scaling of the KSG variance as a function of *N* for the reparameterized log-normal data, cf. Fig. 4(B). While the mutual information estimate on the underlying distribution is severely biased, with our reparameterization we are able to not only return to a regime where we can make unbiased estimates, but also where we have the 1*/N* variance scaling.

**FIG. 5.**
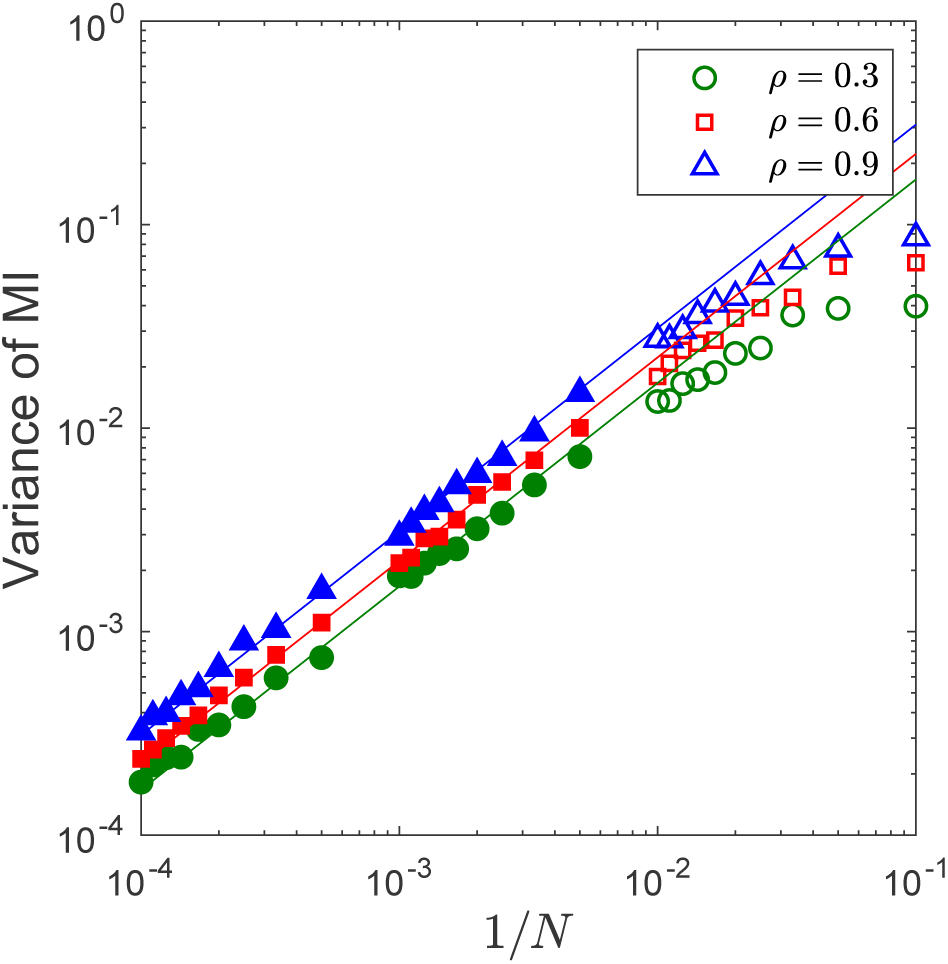
Dependence of the variance of KSG on the sample set size for non-normal data. We repeat the analysis of Fig. 1 for the reparameterized log-normal data of Fig. 4(B), as well as a few other log-correlation coefficients. Here, we have reparameterized from a skewed, heavy tailed distribution, which had biased information estimates. Nonetheless, the scaling 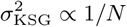 still holds, as illustrated by straight line fits, which have slopes of exactly 1. Empty symbols were not used to fit the straight lines.

A similar reparameterization prescription works for estimating mutual information between higher dimensional variables, although the problems of undersampling are amplified in this case. We first transform each component of the data into a standard normal variable using Eq. (6). We then estimate the estimator variance by performing a linear fit to variances of partitions and then extrapolating to the full data set size. Finally, we check for the *N* -dependent drift for various *k*, and hence choose a good value of *k*, if one exists. Figure 6 shows application of the approach to a 6-dimensional multivariate normal distribution (three dimensions each for *x* and *y*). As in the one-dimensional case, the estimator does not work without reparameterization (not shown), but it performs quite well for the marginally normalized data despite having to deal with more dimensions.

**FIG. 6.**
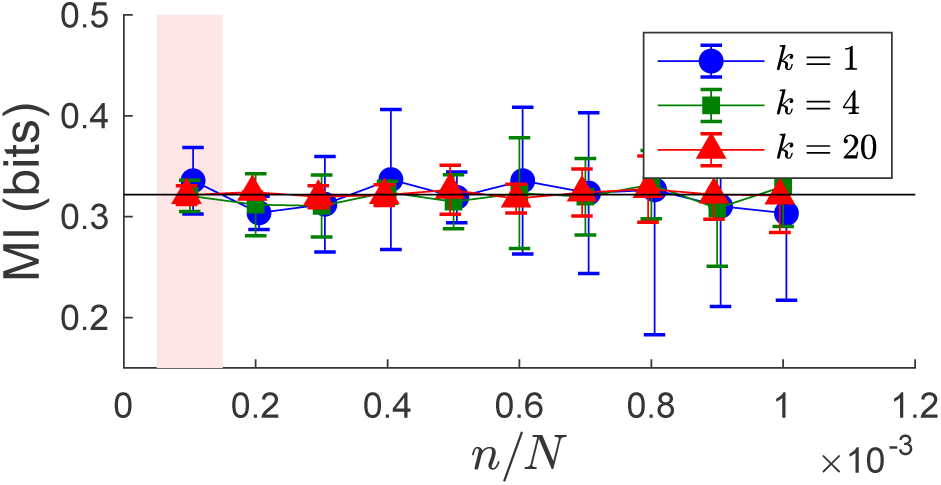
KSG for multivariate data. While other choices could be explored, we chose to start with the same log-normally distributed data as in Fig. 4. We then ro-tate *x* into three components, 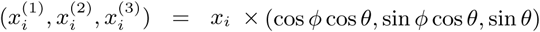, where *ϕ* = *π/*6 and *θ* = *π/*3. We similarly make *y* three dimensional with the same *ϕ* and *θ*. Now KSG needs to find the information between two three-dimensional log-normal variables. For these data, KSG is biased (not shown). However, performing marginal reparam-etereizations for each of the six involved variable components independently, we recover the unbiased performance statistically indistinguishable from Fig. 4: the KSG estimate does not show sample size dependent drift, is consistent for many *k*s, and matches the analytical information value (black horizontal line) for the full data set (pink background).

## III. PRACTICAL GUIDE

MatLab package for performing all of the analyses described above are available from https://github.com/EmoryUniversityTheoreticalBiophysics/ContinuousMIEstimation. In this section, we describe functions in this package, list our specific recommendations for using it to estimate mutual information for continuous variables, and demonstrate how to do so using two experimental data sets.

### A. Functions in the software package

`MIxnyn.m` We distribute the original KSG software (written in C and MatLab) together with our modifications of it. Details for compiling and installing the package are available in the `README` file. This function provides the MatLab interface to the C implementation of KSG. It takes two vectors of samples *x*_*i*_ and *y*_*i*_ as input, where either or both can be multi-dimensional, assumes the usual Euclidean metric on both the *X* and the *Y* space, and produces a single estimate of the mutual information between the two variables.

`findMI_KSG_subsampling.m` This function calculates 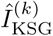 for the full data and its nonoverlapping subsets. It takes two vectors of (potentially multi-dimensional) samples *x*_*i*_ and *y*_*i*_ on the input, as well as a single value of *k* and the vector of *n*, the number of subsets to divide the data into. For each value in the vector *n*, it partitions the data into this many nonoverlapping partitions at random, calculates 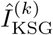 for each subset, and outputs results of all of these calculations. It can additionally make a figure similar to Fig. 3 for a single value of *k*, which allows the user to check for the sample-size dependent drift visually.

`findMI_KSG_stddev.m` This function calculates the variance *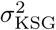* for the full data set, as described above. For this, it takes the output of `findMI_KSG_subsampling.m` (the mutual information values for different subsamples of the data) as well as the data set size *N* as the input. It then calculates the sample variance of *n* values of 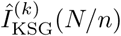 for all available *n* and extrapolates the variance to the full data set size of *N*. If requested, the function can produce a figure similar to Fig. 2, illustrating the procedure and allowing for a visual inspection of whether the variance of subsamples is *?* 1*/N*, as expected.

`findMI_KSG_bias_kN.m` This is the wrapper function that performs our analysis for different values of *k*. It takes the *x*_*i*_ and *y*_*i*_ samples, the list of *k*s to try, and the list of the number of data partitions *n* as the input. It calls the two previous functions sequentially and es-timates 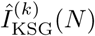 with error bars for every value of *k*. The function can additionally make a figure similar to Fig. 3 for all values of *k* to help find the value *k* for which KSG has the smallest sample size dependent drift and the smallest variance. The function outputs a list of mutual information values with error bars, each corresponding to a specific value of *k*.

`reparamaterize_data.m` The function reparameterizes the data to a standard normal distribution, which, if performed before other estimation steps, should increase the range of applicability of KSG. It takes a vector of samples *x*_*i*_, which must be one dimensional, as the input and returns the reparameterized data as the output.

### B. Application notes

1. Transform each of the components of both *X* and *Y* into the standard normal form using `reparamaterize_data.m.` This should not have any negative effects on the estimation, and may turn out to be extremely advantageous.
2. Do not use bootstrapping and related techniques to estimate variance of the estimator.
3. For a few values of *k*, explore the dependence of the estimates on *k* and the data set size using `findMI_KSG_bias_kN.m` or other functions in the package. Look for a signature of the estimator drift for smaller data set sizes (many partitions), and similarly look for a signature of deviation from ∼ 1*/N* scaling for the variance. These deviations and drift will set the maximum number of data partitions one can explore, and hence will limit the ability to verify whether the estimator is unbiased.
4. Choose the value of *k* for which the estimator shows no statistically significant drift over the largest range of the data set size. If many such *k*s exist, choose the value for which the estimator error bars are the smallest over the range. Note that the estimator should be stable in some range of *k* around the optimal value, but one cannot expect the estimate to be fully independent of *k*.
5. Resist the temptation of subtracting the bias (extrapolating the estimator to *N* → ∞), or declaring the estimator unbiased based only on a small range of *N*. Empirically, about a decade of stability in *N* is needed for this determination. Recall that no estimator is universally unbiased, and so it might be impossible to estimate the information reliably from your data using KSG.
6. If no unbiased *k* is found, try to reduce the dimensionality of your data by any available dimensionality reduction approach. Biases decrease rapidly when the dimensionality decreases. On the other hand, performing any manipulations with data cannot increase the information (by the Data Processing Inequality), and thus one may be able to estimate the lower bound on the true information reliably, with little bias, which may be sufficient for some applications.

### C. Examples

Our software package includes two experimental data sets, showing the utility of the method and allowing one to practice estimation for realistic data.

The first data set comes from the systems biology literature and can be found in `NFkappaBData.mat.` These data were taken with permission from Ref. [23]. The data describe the joint activity of two transcription factors NF-*κ*B and p-ATF-2 measured in 335 individual wild-type mouse fibroblast cells 30 min after exposure to the tumor necrosis factor (TNF) ligand at the concentration of 1.3 ng/mL. The two transcription factors are activated downstream of the same TNF receptor, and hence their activity is correlated. The mutual information between these two sets quantifies this relation. Figure 7 shows application of our method to these data. The figure can be generated by `NFkappaBDataExample.m,` which is included in the distribution.

**FIG. 7.**
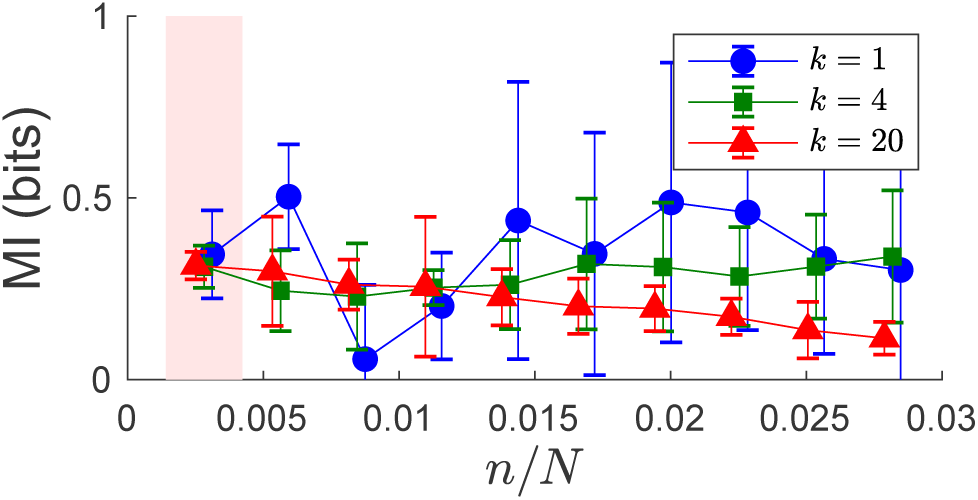
Application of KSG to systems biology data. Mutual between NF-*κ*B and p-ATF-2 activation in mouse fibroblasts 30 min after activation with TNF at 1.3 ng/mL is shown. Data has been marginally reparameterized to a standard normal for this plot (without the reparameterization, estimates are biased). For the full data set size (pink background), standard deviations are extrapolated as detailed above. *k* = 20 shows downwards bias for, at least, large number of partitions. *k* = 1 is unnecessarily noisy. *k* = 4 exhibits a good balance of low drift (bias) and low variance.

The second data set illustrates application of KSG to neurophysiology data and can be found in `BirdSpikingData.mat.` The data have been taken with permission from Ref. [16]. They represent recordings of neural activity from anesthetized Bengalese finches, measured in the motor neurons that control breathing. Here we are analyzing the structure of the spike train itself. The recorded neurons fire only during a particular phase of the breathing cycle, and we are looking at the interspike intervals within such bursts. Specifically, we are estimating the mutual information between two subsequent interspike intervals as one variable, and the following two interspike intervals as the other. Importantly, this is high-dimensional (two dimensions for both *x* and *y*) and non-Gaussian real data. Without reparameterization, questions would remain about the persistent bias of the estimator. However, the marginally reparameterized data in Fig. 8 show no residual bias and a stable estimation for many values of *k* and *N/n*. The figure can be generated by `NFkappaBDataBirdSpikingDataExample.m,` included in the distribution.

**FIG. 8.**
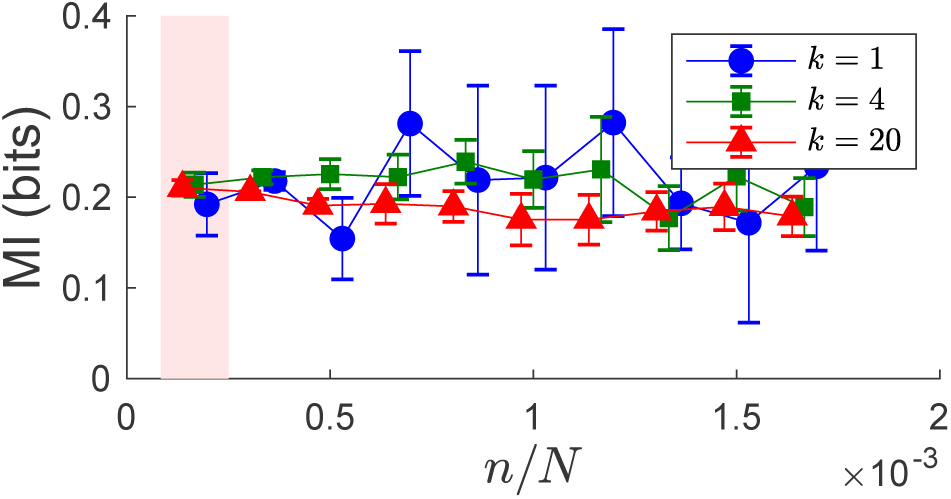
Application of KSG to neurophysiological data. Mutual information between a pair of interspike intervals and the following pair of interspike intervals within a breathing cycle for anesthetized Bengalese finches is being estimated. Despite the high dimensionality and the non-Gaussian nature of the data, we are able to find a stable estimate for the information with 6000 samples. The estimate is stable for many values of *k*, with similar error bars for *k* > 1 (*k* = 1 again gives unnecessarily large error bars). The unreparameterized case (not shown) performs markedly less well.

## IV. DISCUSSION

While mutual information is being used routinely in analysis of modern experimental data sets, high quality, unbiased estimation remains an open problem. In this article, we described our modifications to the well-known Kraskov, Stögbauer, and Grassberger [17] *k* nearest neighbors estimator of mutual information for real-valued data. Our contributions include developing a method for estimating the variance of the estimator, for detecting the presence of bias, and for choosing the optimal value of *k*. Further, we suggest that transforming each marginal data dimension into the standard normal form improves the range of applicability of the estimator, allowing its use even for high-dimensional data sets. We substantiate our choices with extensive numerical investigations. Finally, we provide a MatLab package implementing these modifications to the KSG estimator, as well as a few examples and a practical guide for the workflow. We hope that these developments will be of use to a broad community of physics, quantitative biology, and complex systems researchers.

We end this article with the following observation. As we mentioned in the *Introduction*, there are provably no universally unbiased estimators of mutual information, and thus every estimator—including the one we have developed here—will fail for some data sets. Nothing replaces looking at the data critically and thinking about whether the estimated values make sense and whether there are some patterns in the data that can be used to reduce the dimensionality, to simplify the estimation problem, or to verify the results. Blind application of any algorithm for estimation of mutual information in real-valued data, including application of our modification of the KSG approach, is likely to lead to a failure precisely when the data become interesting.

## ACKNOWLEDGMENTS

We are thankful to Rachel Conn, Sam Sober, and other users of preliminary versions of our software packages for valuable feedback. We thank Raymond Cheong, Kyle Srivastava, Andre Levchenko, and Samuel Sober for providing experimental data for the examples in this work. CMH was supported in part by the Woodruff Scholarship at Emory University and the NSF Center for the Physics of Biological Function (PHY-1734030). IN was supported in part by NIH Grant 1R01NS099375 and NSF Grant IOS-1822677.

## APPENDIX

We are trying to fit a model for the dependence of the KSG estimator variance on the sample size of the form

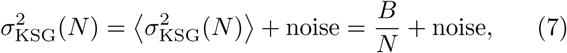

where the angular brackets denote the expectation value. By subsampling or partitioning the data, we can get (noisy) samples of the variance at smaller values *N*_*i*_ than the actual maximum data set size, which we denote *N*. For each of these samples 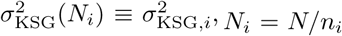, *N*_*i*_ = *N/n*_*i*_, can be evaluated empirically, with *n*_*i*_ being the number of partitions of the data. For example, if we split the data into *n*_*i*_ = 3 parts, we calculate the KSG mutual information for these 3 subsets, and we then estimate the variance at this *N*_*i*_,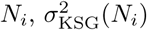 as the empirical variance of the three estimated values. Note that there can be multiple equal values of *n*_*i*_ since data can be partitioned into the same number of parts in many different ways.

The variable 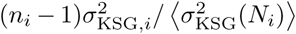 obeys the *χ*^2^ distribution with *n*_*i*_ *-* 1 degrees of freedom, 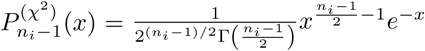. Assuming independence of all 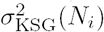 at different values of *i*, and using Eq. (7), we view the product 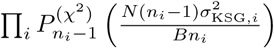 as a likelihood function for *B*. Differentiating w. r. t. *B*, we find the maximum likelihood (ML) solution

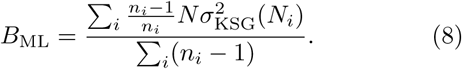

Thus the estimate of the KSG variance at the full data set size *N* is

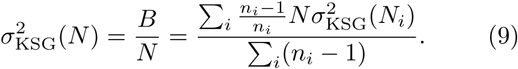

We then calculate the standard error of *B* and, with that, of the variance itself as the inverse of the second derivative of the log-likelihood at the maximum likelihood value:

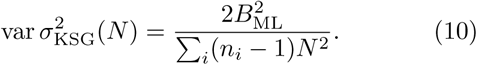

These results are used for estimation of the KSG variance and its error bars in the main text.

